# Self-cutting and integrating CRISPR plasmids (SCIPs) enable targeted genomic integration of genetic payloads for rapid cell engineering

**DOI:** 10.1101/2020.03.25.008276

**Authors:** Darin Bloemberg, Daniela Sosa-Miranda, Tina Nguyen, Risini D. Weeratna, Scott McComb

**Author notes:** Corresponding Author: Dr. Scott McComb, National Research Council Canada, 1200 Montreal Road, Building M54, Ottawa, ON, Canada K1A 0R6.

## Abstract

Since observations that CRISPR nucleases function in mammalian cells, many strategies have been devised to adapt them for genetic engineering. Here, we investigated self-cutting and integrating CRISPR-Cas9 plasmids (SCIPs) as easy-to-use gene editing tools that insert themselves at CRISPR-guided locations. SCIPs demonstrated similar expression kinetics and gene disruption efficiency in mouse (EL4) and human (Jurkat) cells, with stable integration in 3-6% of transfected cells. Clonal sequencing analysis indicated that integrants showed bi- or mono-allelic integration of entire CRISPR plasmids in predictable orientations and with limited indel formation. Interestingly, including longer homology arms (HAs) (500 bp) in varying orientations only modestly increased knock-in efficiency (∼2-fold). Using a SCIP-payload design (SCIPpay) which liberates a promoter-less sequence flanked by HAs thereby requiring perfect homology-directed repair (HDR) for transgene expression, longer HAs resulted in higher integration efficiency and precision of the payload but did not affect integration of the remaining plasmid sequence. As proofs-of-concept, we used SCIPpay to 1) insert a gene fragment encoding tdTomato into the *CD69* locus of Jurkat cells, thereby creating a cell line that reports T cell activation, and 2) insert a chimeric antigen receptor (CAR) gene into the *TRAC* locus. Here, we demonstrate that SCIPs function as simple, efficient, and programmable tools useful for generating gene knock-out/knock-in cell lines and suggest future utility in knock-in site screening/optimization, unbiased off-target site identification, and multiplexed, iterative, and/or library-scale automated genome engineering.

## Introduction

Many systems of stably introducing synthetic genes into mammalian cells exist. The most common, lentiviruses, are mainstays of cell biology and are used for such diverse functions as: expressing novel proteins, stably introducing shRNA, delivering barcoded CRISPR-Cas components, expressing fluorescent markers or tagged proteins, and gene therapy. Despite their widespread use, lentiviruses have several limitations: they require high-level training/facilities and precursor product preparation; their integration location is relatively uncontrollable; their envelope proteins are immunogenic; and their payload size and cell specificity are limited (Broussau et al. 2008; Sakuma et al. 2012; Vannucci et al. 2013).

Since observations that Type II CRISPR endonucleases (i.e. Cas9) function in mammalian cells, many strategies have been devised that exploit their reprogrammable capacity for inducing double-strand breaks (DSBs) for performing gene editing. This includes delivering donor sequences via AdV/AAV, high-yield PCR product, plasmid, or single stranded DNA oligonucleotides, and CRISPR-Cas components via plasmid, RNA, RNP, or replication-incompetent viruses (Jin et al. 2019; Ehrke-Schulz et al. 2017; Mangeot et al. 2019; Liu et al. 2018; Gao et al. 2018; Moore et al. 2015; Tálas et al. 2017; Sawatsubashi et al. 2018; Leonetti et al. 2016; Zhu et al. 2015; Lundin et al. 2020; Maresca et al. 2013; Schmid-Burgk et al. 2016). In addition to targeted gene editing for fixing germ-line mutation diseases, the current “killer app” for inserting synthetic transgenes is CAR-T, whereby T cells are programmed to target cancer cells using an engineered chimeric antigen receptor (CAR) (Bloemberg et al. 2019). Of note, several non-lentiviral methods have shown practical use for CAR-T engineering. The Sadelain group inserted a chimeric antigen receptor (CAR) transgene into the *TRAC* locus by electroporating T cells with Cas9/gRNA RNPs and delivering the CAR transgene using rAAV6 (Eyquem et al. 2017). The Chen group also recently generated CAR-T cells by electroporating Cpf1 mRNA while delivering gRNA and donor DNA sequences via AAV, a method which enabled genetic disruption of a second location (Dai et al. 2019). The Marson lab similarly delivered Cas9/gRNA RNP into T cells and used high-output PCR products as donor templates to correct a pathological *IL2RA* defect and replace endogenous TCR genes with a cancer antigen-specific transgenic TCR (Roth et al. 2018).

Aside from methods intended to manufacture CAR-T cells at large scales, several simpler nuclease-based systems have shown practical use. Using a canine hemophilia genetic model, transfecting a single plasmid (or transducing single AdV) containing a Tet-ON Cas9 cassette and a donor DNA template flanked by gRNA bait sites into cultured cells enabled inserting this donor repair sequence at the *cFIX* mutation site, thereby generating functional protein (Gao et al. 2018). Utilizing only DNA plasmids, introducing specific ZFNs along with a donor plasmid containing oppositional recognition sites around a DNA payload permitted integration of genes up to 15 kb in the *AAVS1* locus (Maresca et al. 2013), a strategy coined ObLiGaRe which has recently been extended to stably integrate an inducible Cas9 cassette into *AAVS/Rosa1* (Lundin et al. 2020). Similarly, utilizing HEK cells with stable Cas9 expression and up to three separate DNA plasmids, a strategy of NHEJ-mediated integration of plasmid-derived DNA sequences was engineered with modular frame control, allowing payload integration at CRISPR-guided locations and gene-specific transgene expression (Schmid-Burgk et al. 2016).

Inspired by lentiviral vectors and these other plasmid-based gene delivery systems, we developed a method for targeted integration of DNA payloads at specific genomic sites by engineering CRISPR-Cas9 plasmids to simultaneously cut genomic sequences and themselves: a strategy we call self-cutting and integrating plasmids, SCIP. Here, we demonstrate that SCIPs represent easy-to-use genetic engineering tools for generating knock-out cell lines, fluorescent reporter knock-in cell lines, and inserting DNA payloads at specific genomic sites. As SCIPs are easily-reprogrammable and single-component genetic manipulation tools, we anticipate their public availability will enable widespread use in routine cell line engineering, serial/repetitive gene manipulation, genome editing target optimization, and multiplexed and/or library-scale genome editing.

## Results

### Self-cutting and integrating plasmids (SCIP) as a non-viral strategy for mammalian genome editing

We hypothesized that including a “bait” site for the corresponding gRNA in a CRISPR-Cas9 plasmid could generate an easy-to-modify and -manufacture single-component gene editing tool (Fig. 1A). Upon transfection into mammalian cells, this plasmid would produce double-strand breaks (DSBs) and self-linearize, possibly leading to its integration (Fig. 1A). This ability to stably integrate user-defined DNA payloads would mimic the function of lentiviral transgene delivery; however, we hypothesized that SCIPs could integrate at CRISPR-guided locations (Fig. 1A).

**Figure 1.**
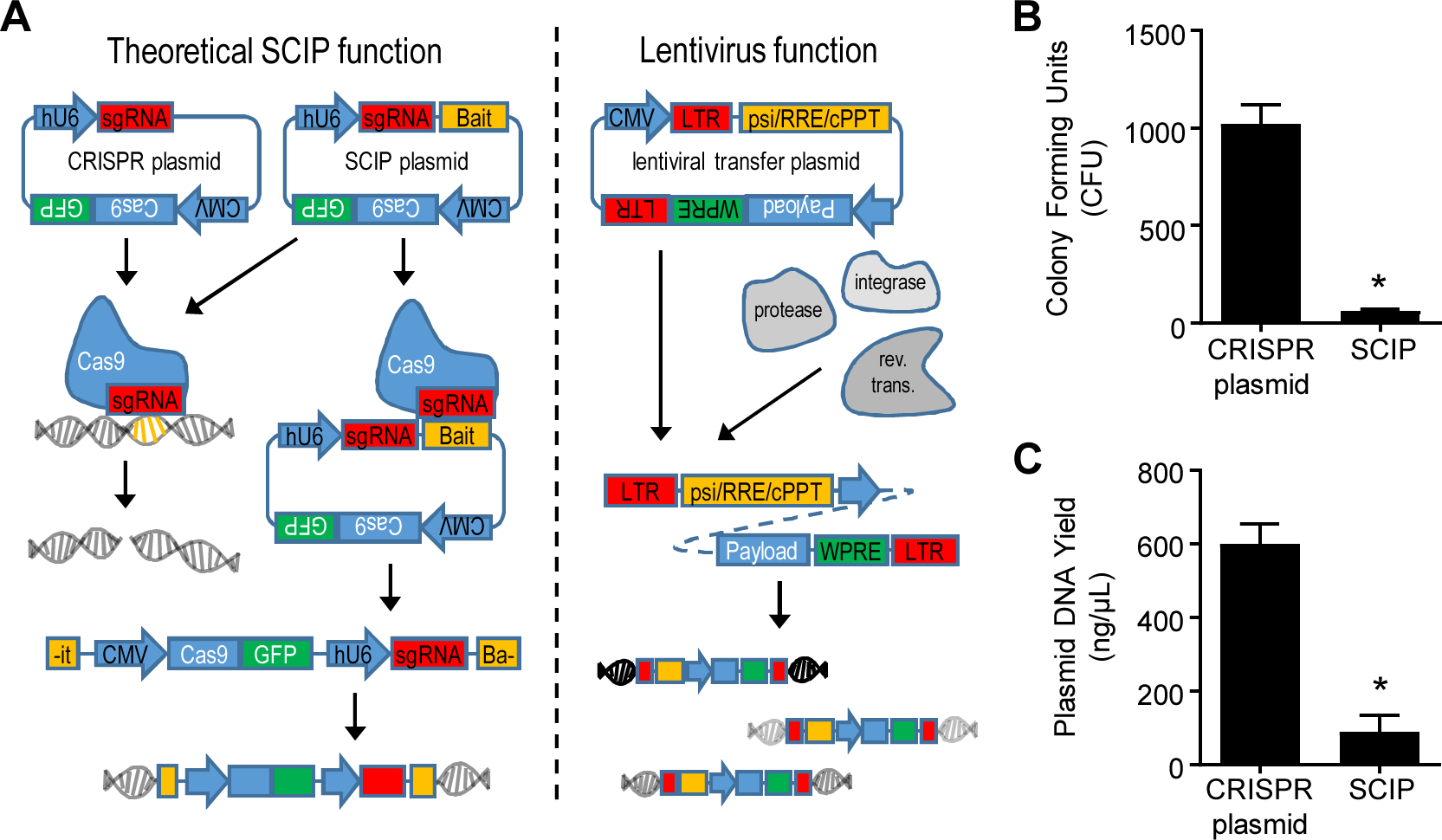
Self-cutting and integrating plasmid (SCIP) function and lack of bacterial stability. (A) Overview of theoretical SCIP function. Identification and cleavage of the bait site by the corresponding gRNA-Cas9 complexes produced within mammalian cells linearizes the plasmid while performing a genomic DSB. In this way, a single, modular plasmid allows stable integration of user-defined DNA payloads, in a manner similar to lentiviruses. (B & C) Colony forming units (B) and mini-prep DNA yield (C) in SCIP compared to guide-containing CRISPR plasmids. Results in (B) and (C) represent means +/- SEM of 3 independent experiments. Asterisk (*) represents significant difference, as calculated using T-tests (p<0.05).

We created a SCIP cloning plasmid by adding a second “Golden Gate” site to the popular GFP-expressing CRISPR-Cas9 plasmid px458 (see Methods for details) to be used for inserting bait sequences (pQC; Supplemental Fig. S1A). However, regardless of the specific guide+bait sequences used (Supplemental Table S1), self-cutting plasmids proved initially very inefficient to assemble and produce, as successful ampicillin-resistant transformant colony numbers were reduced (p<0.05) by 95% (Fig. 1B) and colonies which did survive produced 90% less (p<0.05) plasmid DNA (Fig. 1C).

### Inserting an intron into the Cas9 sequence significantly improves self-cutting plasmid stability and production in bacteria

We theorized this instability was due to leaky Cas9 and gRNA production in DH5α *E. coli* cells used for molecular cloning and plasmid production, which would cause plasmid degradation (Fig. 2A). To prevent processing of these transcripts in bacteria cells, we inserted a mammalian intron into the Cas9 plasmid genetic sequence, a strategy previously used to increase the production of cleavable CRISPR components (Petris et al. 2017) (we termed this plasmid quick-change with intron, pQCi; Fig. 2B and Supplemental Fig. S1B). Plasmids targeting three different sites of human *TRAC* were created with guide sequences, bait sequences, or both, in pQC and pQCi (generating the 8 plasmids in Supplemental Fig. S1B). Not only were transformant colony counts (Fig. 2C), plasmid DNA yield (Fig. 2D), and qualitative DNA quality (Fig. 2E) unaffected (p>0.05) by the intron, but guide+bait plasmid instability was completely reversed in pQCi (Fig. 2C & 2D). This improved stability of guide+bait-containing plasmids was also observed in Stbl3 and BL21 cell lines (Fig. 2F & 2G), indicating that the phenomenon of leaky bacterial Cas9/gRNA expression is not restricted to DH5α.

**Figure 2.**
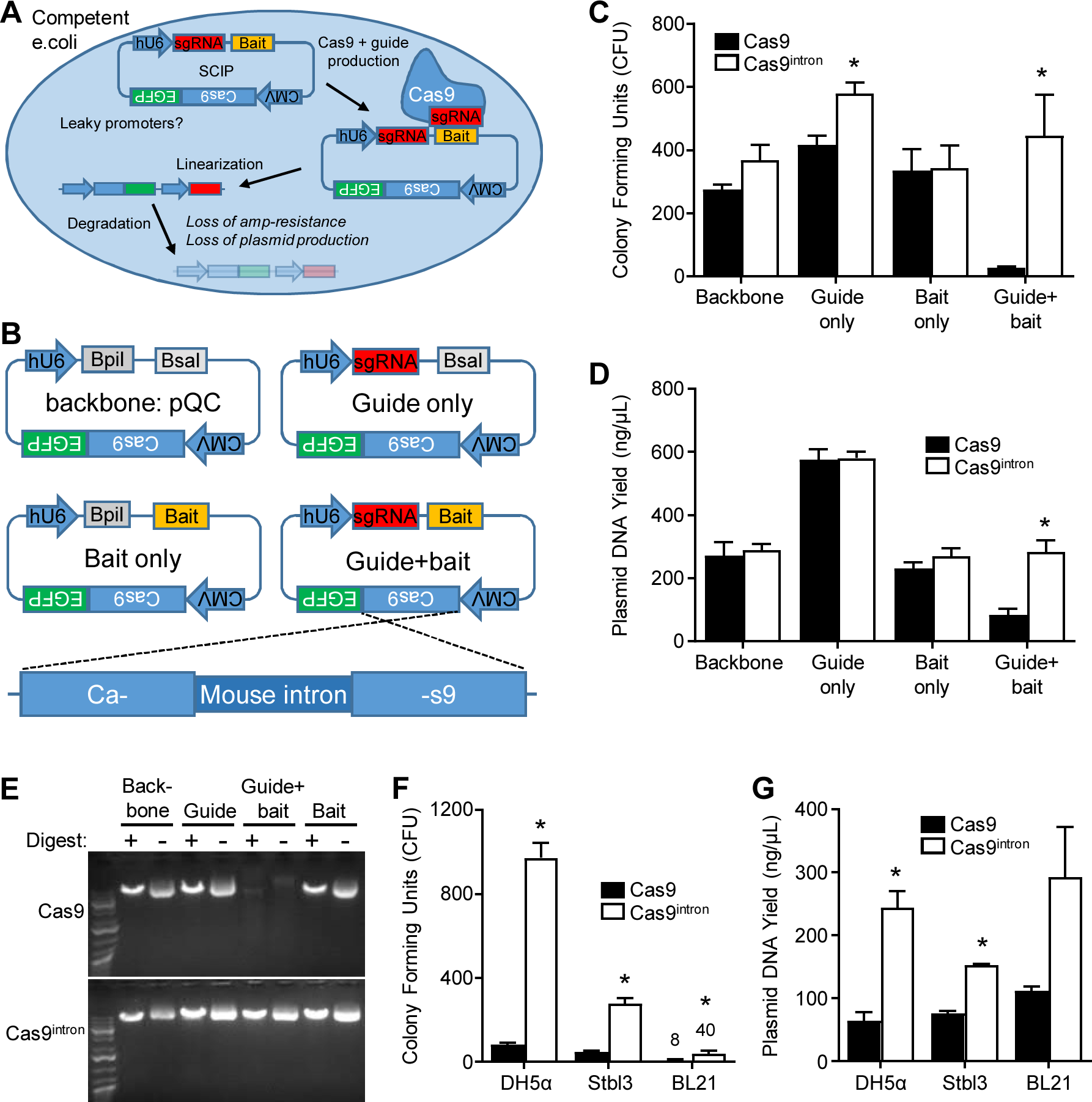
Inserting an intron into the Cas9 sequence significantly improves SCIP stability and production in bacteria. (A) Proposed mechanism contributing to reduced SCIP stability in bacteria cells. Leaky CMV and hU6 promoters lead to Cas9/gRNA complex production and subsequent plasmid cleavage. (B) Overview of SCIP backbone (pQC), plasmid, and intron structures. See Supplementary Fig. S1 for an overview of these plasmids. (C) Number of colonies formed from cloning reactions when creating various normal/Cas9 and intron-containing plasmids. (D) Plasmid DNA production from various normal/Cas9 and intron-containing plasmids. (E) Plasmid DNA quality as demonstrated in agarose gel. (F & G) Colony counts (F) and plasmid DNA yield (G) when using the indicated competent cell strains to generate guide+bait plasmids. Results in (C), (D), (E), and (F) represent means +/- SEM of 4-5 independent experiments. Asterisks (*) represent significant difference between Cas9 and Cas9^intron^, as calculated using T-tests (p<0.05).

### Self-cutting plasmids display similar expression and gene disruption efficiency as CRISPR-Cas9 plasmids

Given the potential for self-cutting to destabilize plasmid function in mammalian cells we next tested the function and stability of self-cutting plasmids by electroporating mouse EL4 cells with various plasmid formats. Here, GFP expression was similar (p>0.05) in pQC and pQCi empty, guide, and bait plasmids, demonstrating that processing was unaffected by the intron (Fig. 3A – 3C). Importantly, GFP expression produced by pQCi-guide+bait was higher (p<0.05) compared to pQC and was similar to empty, guide, and bait counterparts (Fig. 3A – 3C). When evaluating gene disruption efficiency for plasmids targeting *B2m* or *Pdcd1*, px458-guide, pQCi-guide, and pQCi-guide+bait produced similar percentages (p>0.05) of knock-out cells (Fig. 3D & 3E), again suggesting that the intron does not interfere with Cas9 and that self-cutting plasmids remain functional. Thus, restored stability of intron-containing self-cutting plasmids in bacteria cells had apparently normal function in mammalian cells warranting further investigation of their integration activity.

**Figure 3.**
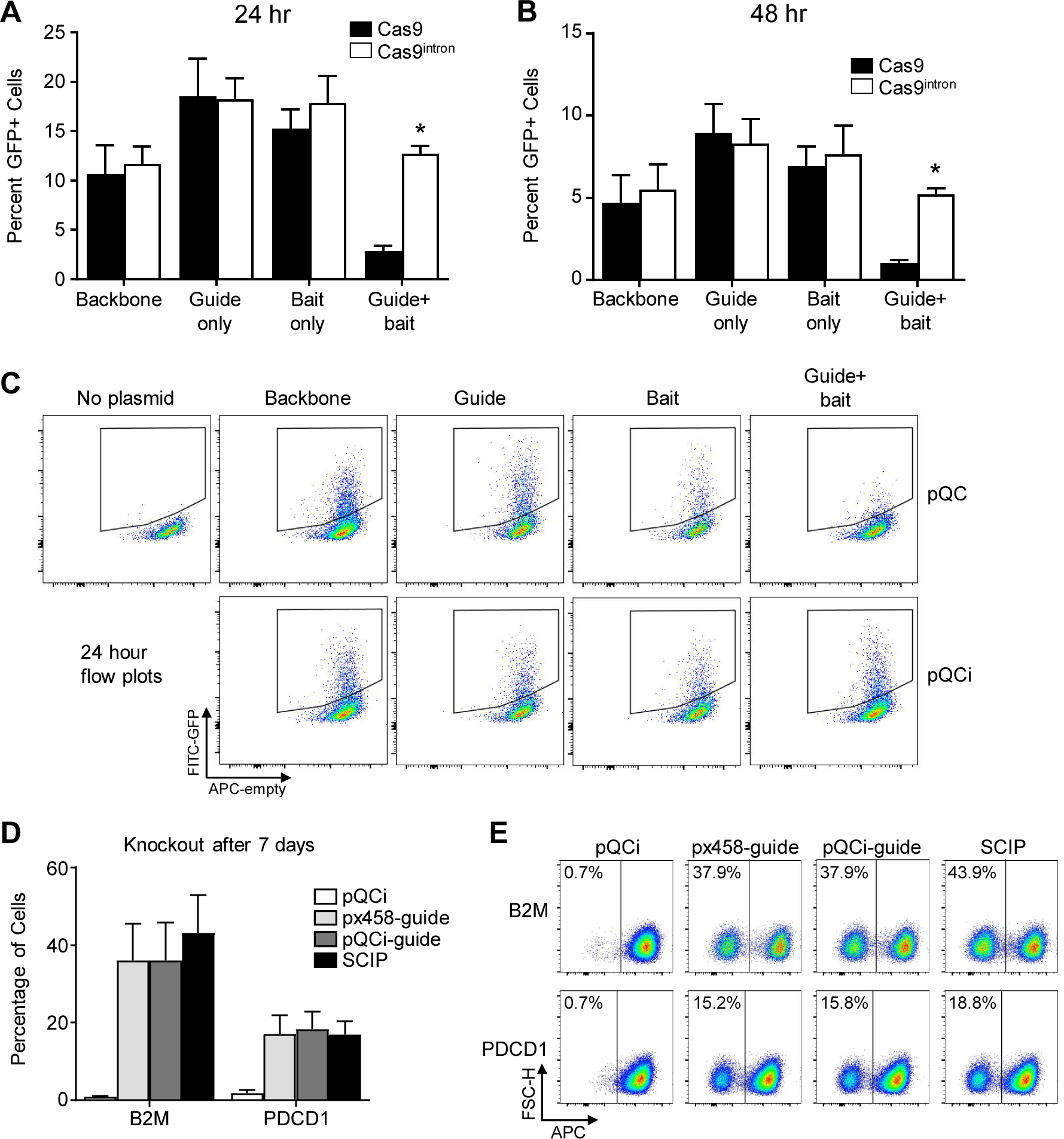
SCIPs display similar expression and gene disruption efficiency as CRISPR-Cas9 plasmids. (A – C) The indicated plasmids were electroporated into mouse EL4 cells and assessed for GFP production using flow cytometry 24- (A & C) and 48-hours (B) after transfection. (D & E) Cells were similarly electroporated with the indicated plasmids containing gRNA targeting *B2m* or *Pdcd1* and analyzed for expression of the respective protein 7 days later. Results in (A), (B), and (D) represent means +/- SEM of 3 independent experiments. Asterisks (*) represent significant difference between Cas9 and Cas9^intron^, as calculated using T-tests (p<0.05).

### SCIPs integrate into CRISPR-guided locations

SCIP integration should result in stable expression of all plasmid ORFs. Here, this includes CMV-driven Cas9-eGFP, enabling simple knock-in detection using flow cytometry (Fig. 4A). Using the same plasmids constructed targeting mouse *B2m* and *Pdcd1*, the integration efficiency (the percentage of knock-out cells that are GFP-positive) increased (p<0.05) from <0.5% to 3% for *B2m* and from 2% to 6% for *Pdcd1* when EL4 cells were electroporated with SCIPs as compared to guide-only px458 and pQCi (Fig. 4A & 4B). Similarly, the integration precision (the percentage of GFP-positive cells that are knock-outs) increased (p<0.05) from 50% to 80% for *B2m* and from 45% to 60% for *Pdcd1* with SCIPs compared to guide-only plasmids (Fig. 4A & 4B). Importantly, these knock-out/knock-in events were stable and unchanging, as profiles 7 days post-transfection (Fig. 4) remained after 7 additional days (Supplemental Fig. S3), and much longer for clonal experiments discussed below. Therefore, self-linearization enabled at least a 5-fold increase in plasmid integration and 40% increase in the targeted precision of integration events.

**Figure 4.**
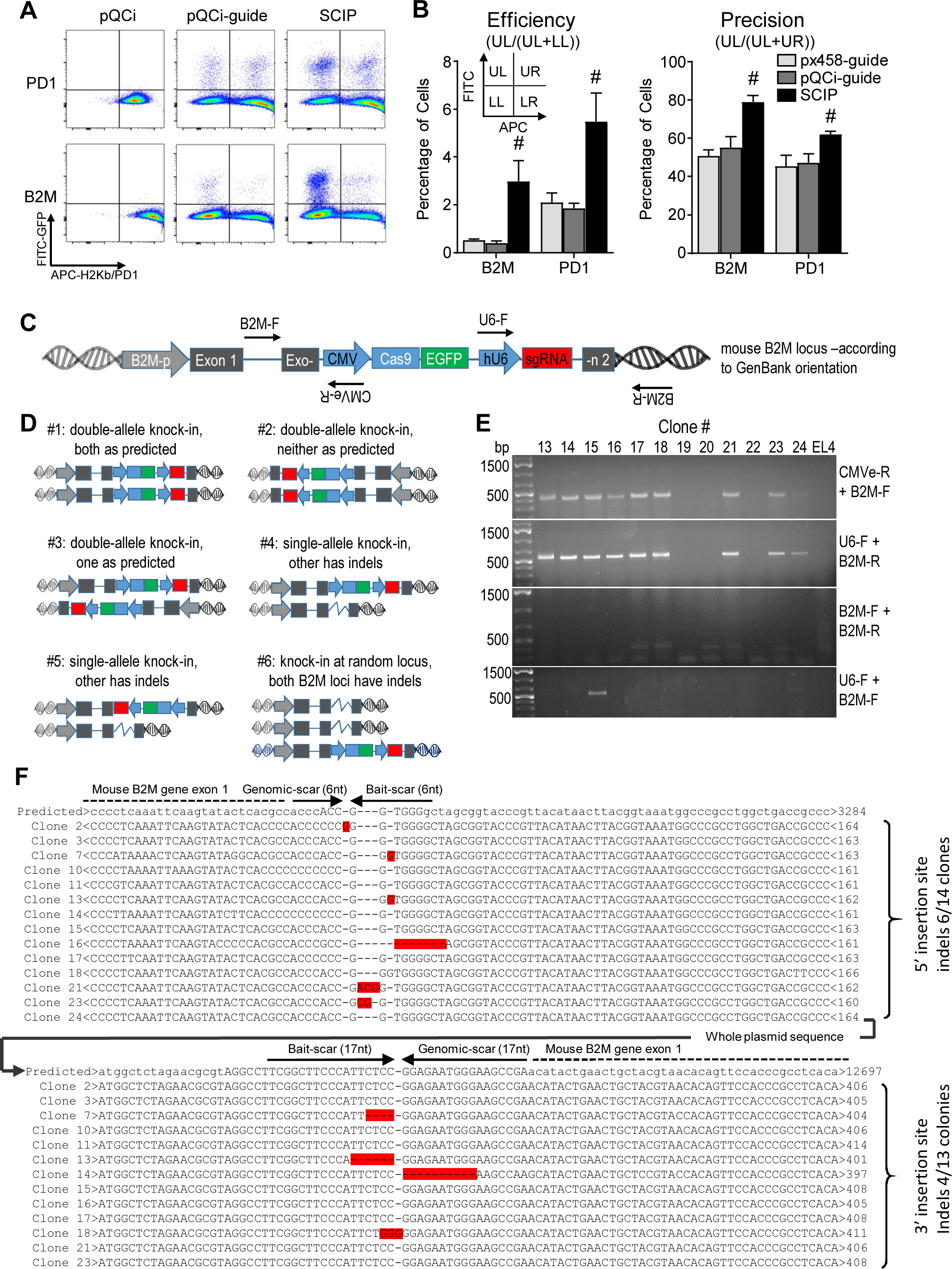
SCIPs integrate into CRISPR-guided locations. (A & B) EL4 cells were electroporated with the indicated plasmids and assessed for gene knock-out and SCIP knock-in 7 days later. (A) FITC versus APC-[marker] flow cytometry plots allow evaluation of knock-out and knock-in events, quantified as efficiency and precision percentages (B), as defined in Figure panel insets. (C) EL4 cells electroporated with *B2m*-targeted SCIP were single-cell cloned and analyzed for knock-in configurations using the indicated PCR primers. (D) Illustration of possible SCIP integration configurations. (E) PCR DNA bands resulting from the indicated primers from clones #13-24. (F) Sanger sequencing results from measurable PCR reactions (*B2m*-F + CMVe-R or U6-F + *B2m*-R) produced from clones showing configuration #1 in (D). Results in (B) represent means +/- SEM of 4 independent experiments. Pound signs (#) represent significant difference between the indicated group and all other groups, as calculated using 1-way ANOVA (p<0.05).

We hypothesized that SCIP integration should favour a specific orientation to prevent re-forming gRNA target sites (Supplemental Fig. S2B). To validate the presence and directionality of SCIP integration, EL4 cells were electroporated with *B2m*-targeted SCIPs and those displaying both deficiency for *B2m* expression and stable GFP expression were single-cell sorted one week later. Genomic DNA extracted from these clonal cultures was then assessed using the PCR strategy outlined in Fig. 4C and Supplemental Table S2. In addition to the predicted direction, 5 other possibilities causing simultaneous knock-out and knock-in are illustrated (Fig. 4D). Of 36 measurable clones, 14 (39%, examples #13, 14, 16, 24 in Fig. 4E) showed double-allele knock-in and 12 (33%, examples #17, 18, 21, 23 in Fig. 4E) showed single-allele knock-in with presumed CRISPR-Cas9-induced insertion or deletion mutations (indels) likely present in the other allele (Table 1). An additional 5 clones (14%, example #20 in Fig. 4E) displayed wild-type bands and no *B2m*-specific integration and were therefore categorized as double-allele indels with assumed knock-in at a random location. Of note, these findings are based on cell surface protein expression and PCR bands and we did not confirm the genuine presence of indels here.

**Table 1:**
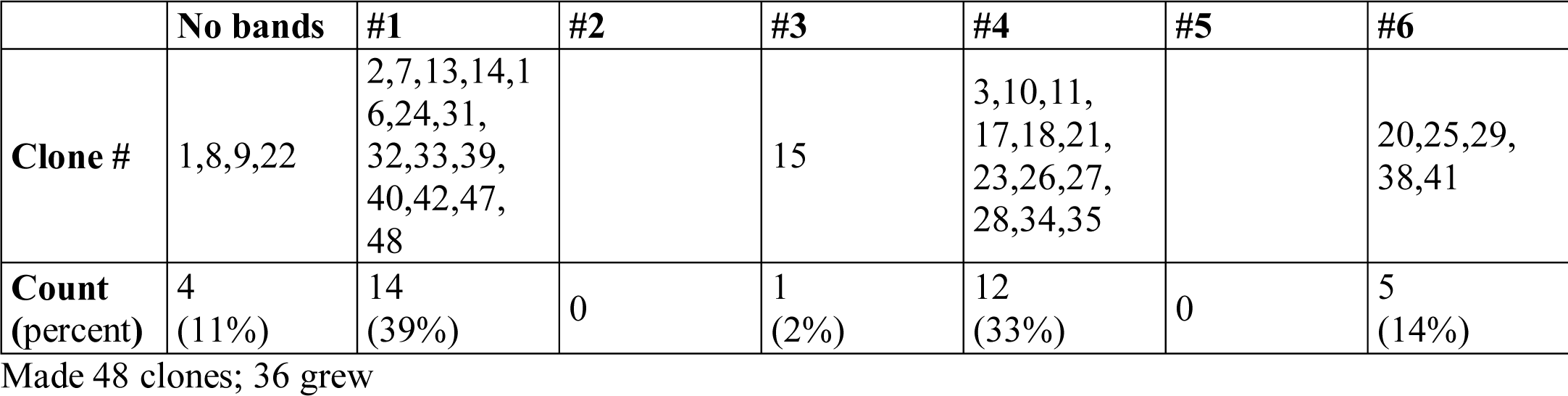
EL4-*B2m*-SCIP Genomic PCR Results.

|  | No bands | #1 | #2 | #3 | #4 | #5 | #6 |
| --- | --- | --- | --- | --- | --- | --- | --- |
| **Clone #** | 1,8,9,22 | 2,7,13,14,16,24,31,32,33,39,40,42,47,48 |  | 15 | 3,10,11,17,18,21,23,26,27,28,34,35 |  | 20,25,29,38,41 |
| **Count (percent)** | 4 (11%) | 14 (39%) | 0 | 1 (2%) | 12 (33%) | 0 | 5 (14%) |

Made 48 clones; 36 grew

PCR products from the 14 clones showing double-allele knock-in were sequenced to confirm the integration mechanism (Figure 4F). Here, the side with 17 bp homology displayed more consistent ligation than the side with 6 bp homology (4 versus 6 clones with indels). However, the inverted homologous regions remained in all clones, providing important insight regarding the mechanism of these repair events (Figure 4F). Overall, this shows that SCIPs enable targeted genomic integration of large genetic payloads (a whole plasmid of ∼9400 bp in this case) at efficiencies feasible for cell engineering.

### Homology arm length and orientation minimally affect SCIP integration efficiency and precision

As SCIP integration appeared directional, we next constructed plasmids with increasing homology arm (HA) lengths surrounding the mouse *B2m* CRISPR site, expecting this might improve knock-in efficiency and precision. Here, instead of differently-sized micro-HAs (17 and 6 bp), HA length was designed to be equal on each side of the plasmid bait site (Fig. 5A, Supplemental Fig. S1C). These plasmids were generated in two formats: 1) inverted homology, in a similar orientation to that present in bait-containing plasmids and 2) forward homology, in an overlapping orientation similar to typical DNA payloads designed for HDR after plasmid linearization (Fig. 5A).

**Figure 5.**
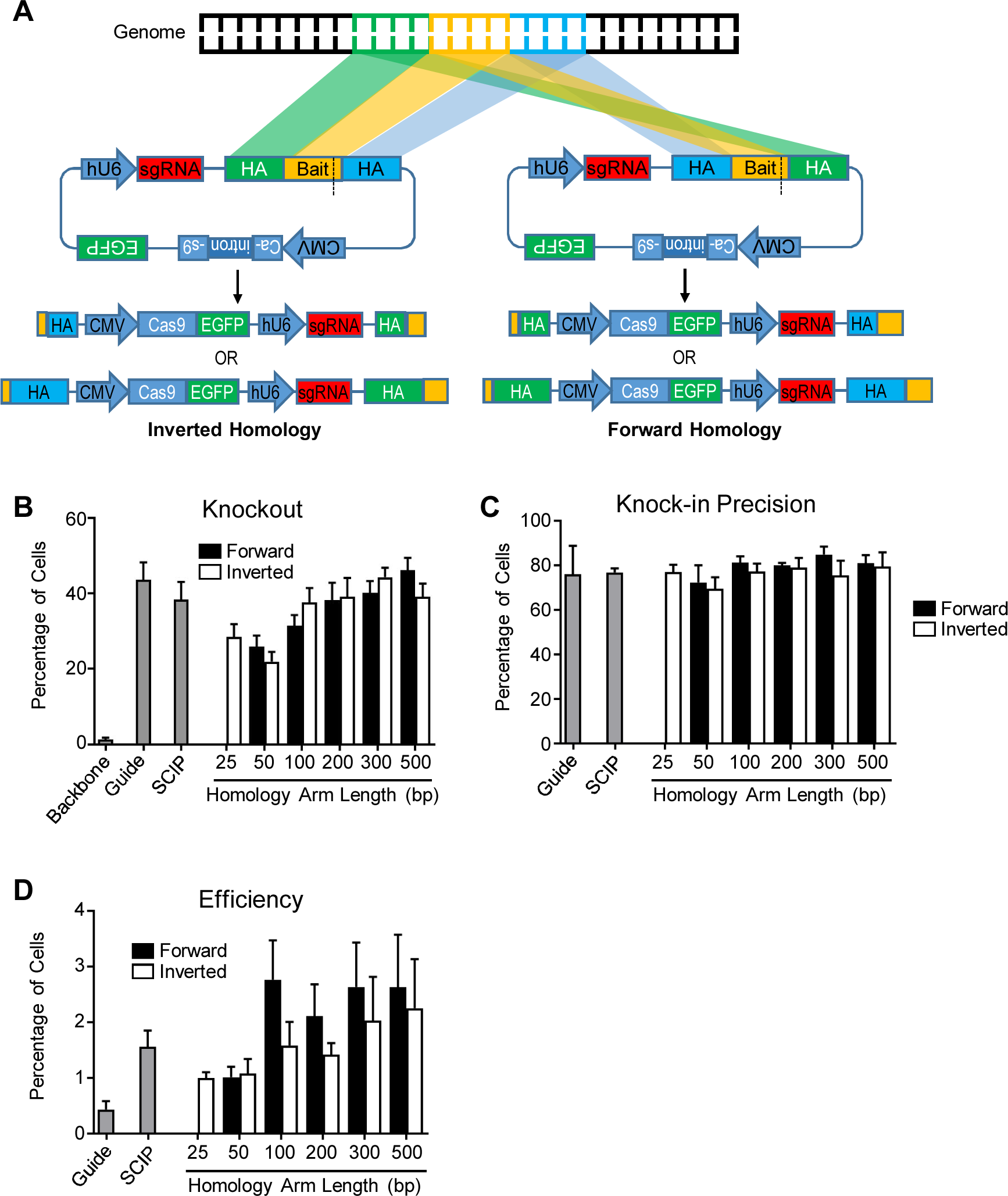
Homology arm length and orientation minimally affect SCIP integration efficiency and precision. (A) Overview of structural design for SCIPs with longer HAs. (B – D) EL4 cells were electroporated with SCIPs targeting *B2m* and assessed for knock-out/knock-in 7 days later. (B) Percentage of *B2m* knock-out cells and corresponding precision (C) and efficiency (D) scores. Results in (B), (C), and (D) represent means +/- SEM of 4 independent experiments.

While there was a general trend for increasing *B2m* knock-out efficiency in EL4 cells with increasing HA length, there was no significant (p>0.05) difference compared to pQCi-guide and SCIP and no apparent effect of HA orientation (Fig. 5B). Similarly, HA length and orientation did not affect knock-in precision, suggesting that 80% is the highest possible with this gRNA (Fig. 5C). Similarly knock-in efficiency showed a trend towards increase along with HA length, but did not significantly affect knock-in efficiency compared to bait-only SCIP, increasing only from ∼1.5 to 3% with 500 bp homology arms (Fig. 5D). These results underline the quite high level of knock-in that can be achieved with the limited homology provide by SCIP bait sites.

### SCIPpay design permits CRISPR-guided integration of specific donor DNA sequences via HDR

Our ultimate interest is enabling easier and higher-throughput directed gene editing, where user-defined DNA payloads can be inserted at specific genomic loci or used to replace endogenous sequences. Therefore, we modified SCIP to contain a second bait site flanking a transgenic DNA payload with HAs (SCIPpay). We hypothesized that such a plasmid could deliver DNA sequences via homology directed repair (HDR) without integrating the remaining plasmid (Fig. 6A, Supplemental Fig.S1D).

**Figure 6.**
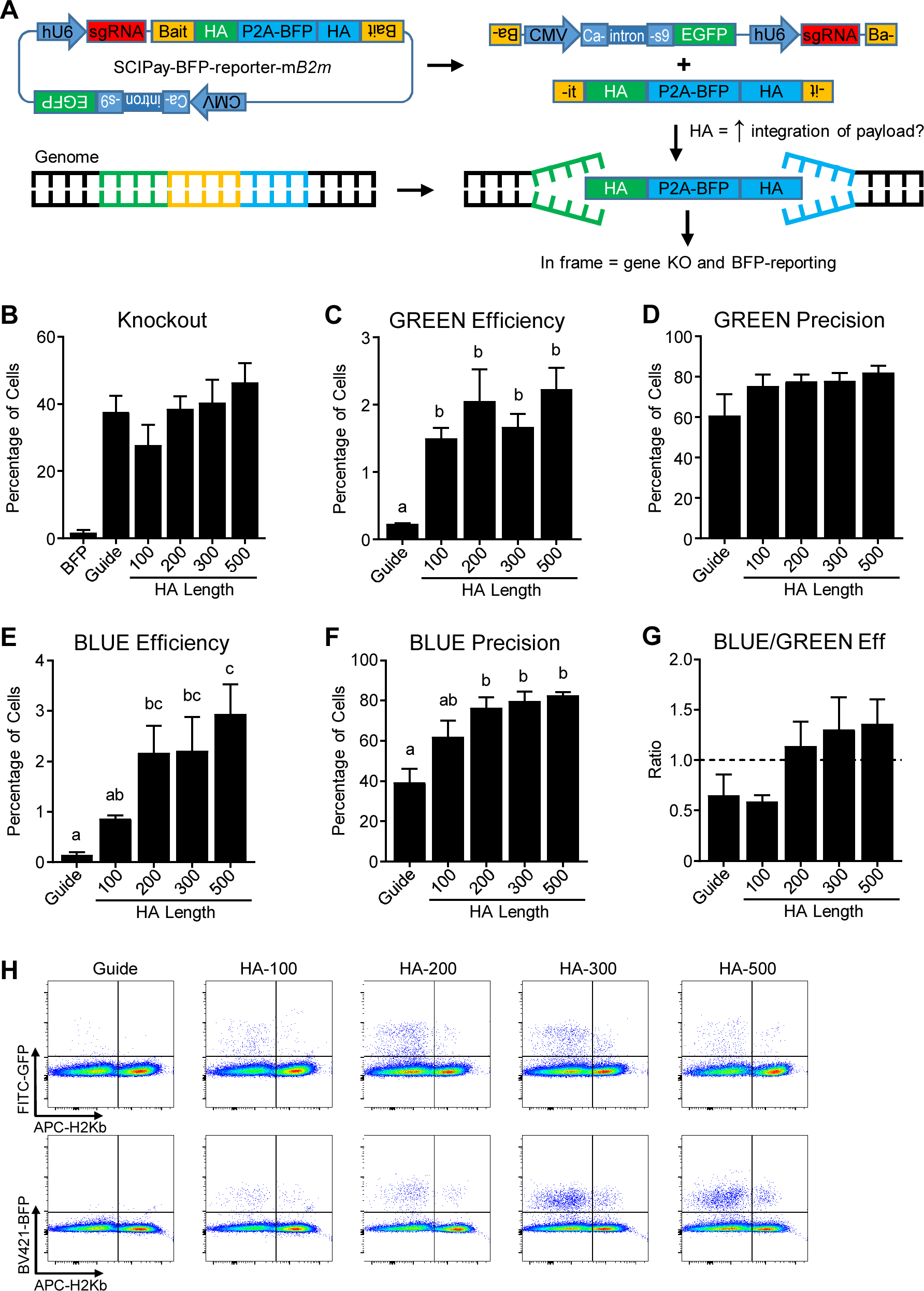
SCIPpay design permits CRISPR-guided integration of specific donor DNA sequences via homology directed repair. (A) Overview of SCIPpay design. Here, HA present in the payload sequence lead to its genomic integration via HDR. (B – G) EL4 cells were electroporated with SCIPpay containing a BFP transgene targeting *B2m* with various HA lengths. (B) Percentage of knock-out cells. (C) Efficiency and (D) precision scores for GFP, denoting integration of the non-payload plasmid sequence. (E) Efficiency and (F) precision scores for BFP, denoting HDR integration of BFP. (G) Ratio of BFP- to GFP-positive cells. (H) Representative flow cytometry pltos. Results in (B – G) represent means +/- SEM of 4 independent experiments. Data was analyzed using 1-way ANOVA and significant differences are indicated with lower case letters, where groups with different letters are significantly different than each other (p<0.05).

To enable this, we first created a fluorescent reporter SCIPpay plasmid containing a promoter-less P2A-BFP transgene, wherein in-frame HDR-mediated transgene integration would result in BFP production (Supplemental Fig. S1D). This construct was given appropriate restriction sites enabling simultaneous insertion of both HA and bait sequences during a single cloning reaction. Continuing with the high-efficiency mouse *B2m* gRNA, SCIPpays were created with different HA length (100, 200, 300, or 500 bp) in-frame with the *B2m* genome flanked by bait sites that yield 6 bp scars facing the payload (Fig. 6A). SCIPpays with HAs produced similar (p>0.05) *B2m* knock-out cells as gRNA-only plasmids (Fig. 6B & 6H). While integration of the Cas9-eGFP portion occurred in 2% of cells transfected with HA-containing SCIPpay, the efficiency and precision of these insertions remained consistent (p>0.05) with increasing HA length (Fig. 6C & 6D). In contrast, the percentage of BFP-positive cells increased with longer HAs, leading to elevated (p<0.05) efficiency and precision (Fig. 6E & 6F). The ratio of BFP-to GFP-positive cells with *B2m* knock-out approached 1.5 with 500 bp HA, indicating that HDR-mediated editing involving the payload sequence occurred 50% more often than any NHEJ editing event for the remaining plasmid, where GFP expression is driven by a constitutive promoter (Fig. 6G). These results demonstrate that SCIPs can be effectively engineered to integrate either whole plasmid or a payload DNA sequence with no definite requirement for large homology arm sequences.

### Using SCIPpay to generate a cell line that reports T cell activation

This HDR efficiency is sufficiently high to perform useful genetic cellular engineering. To demonstrate SCIPpay’s utility in this regard and generate a useful tool for our ongoing CAR development efforts (Bloemberg et al. 2020), we designed a SCIPpay containing a P2A-tdTomato with poly-A sequence transgene (total transgene size = 1815 bp) targeted to human *CD69* with homology arms of ∼500bp (Fig. 7A, Supplemental Fig. S1E).

**Figure 7.**
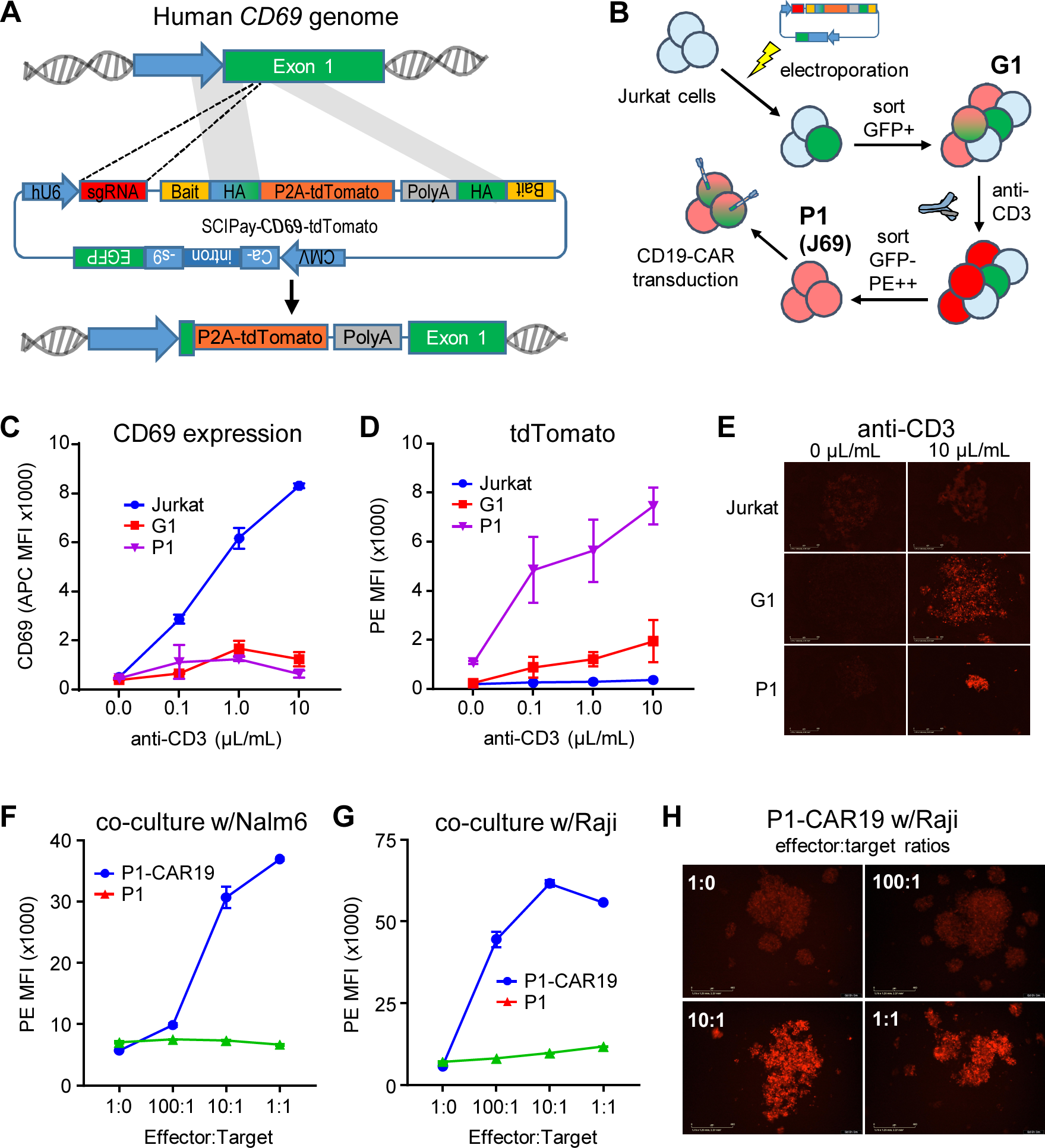
Using SCIPpay to generate a cell line that reports T cell activation. (A) Overview of SCIPpay designed to insert tdTomato into human *CD69*. (B) Outline of cell engineering strategy. After electroporation, successfully transfected cells were sorted, activated, and sorted again to enrich PE-responding cells. (C – E) The indicated cell populations were incubated with increasing anti- CD3 and assessed for surface CD69 (C) and tdTomato (D) expression using flow cytometry. (E) Microscopy images of similar cells. (F – H) P1/J69 was transduced with an anti-CD19 CAR and cultured alongside CD19-positive cells. PE response to Nalm6 (F) and Raji (G & H) co-cultures. Results in (C – H) represent means +/- SEM of 3 independent experiments.

Jurkat human T cells were electroporated with this plasmid, stimulated with anti-CD3 antibody and sorted for GFP and tdTomato expression (Fig. 7B). While simultaneous sorting of tdTomato and GFP makes it difficult to estimate the raw knock-in efficiency, 19% of sorted cells showed tdTomato and 13% of cells showed both tdTomato and GFP expression in CD3-stimulated G1 sorted cells after 8 days (Supplemental Fig. S4A). G1 sorted cells were then re-stimulated with CD3/CD28 beads and re-sorted according to Fig. 7B, producing a polyclonal population (P1 or J69) with medium-low basal PE/tdTomato expression in 100% of cells. When incubated with anti-CD3 antibody to induce activation, normal Jurkat cells displayed a dose-response increase in surface CD69 expression as assessed using flow cytometry, while intermediate (G1) and final reporter (P1) populations showed low CD69 expression (Fig. 7C). Conversely, G1 and P1 displayed a dose-response increase in PE/tdTomato with anti-CD3 administration that mimicked the CD69 response observed in normal Jurkat cells (Fig. 7D & 7E), demonstrating the ability to properly report T cell activation.

While polyclonal J69 produced consistent data, we derived clonal cell lines hypothesizing this would further reduce variability. Here, 24 single-cell clones were isolated from the polyclonal population and assessed for responsiveness (Supplemental Fig. S4). While most clones (20 of 24) functioned like the polyclonal population (CD69 negative and increased PE upon anti-CD3/CD28 stimulation), 3 clones also displayed CD3-dependent GFP expression responses and tdTomato raw signal magnitudes varied significantly (Supplemental Fig. S4). These findings highlight the need for robust screening strategies and emphasize the promiscuity and randomness of using Cas9 for genome editing.

The utility of J69 was further investigated by transducing them with a CD19-targeted CAR lentivirus (Bloemberg et al. 2020) and assessing their ability to report CAR-mediated activation (Fig. 7B). Here, J69-CAR19 cells increased PE/tdTomato expression at increasing target cell ratios when cultured alongside CD19-positive Nalm6 (Fig. 7F) or Raji (Fig. 7G & 7H) leukemia cells, responses that were not observed in non-transduced J69.

### Using SCIPpay to insert a CAR into the T cell receptor alpha constant (*TRAC*) gene

As a final proof-of-concept, a SCIPpay was designed containing an anti-EGFR CAR with a P2A-BFP transgene (total transgene size = 2043bp) targeted to human *TRAC* (Fig. 8A). Here, delivering a single plasmid causes TCR-knock-out and stable CAR integration into a specific genetic site in HDR-edited cells which are indicated by BFP. As before, this plasmid’s function was demonstrated by electroporating and sorting Jurkat cells (Fig. 8B). In this experiment, BFP expression was minimal 3 days post-electroporation (0.35%), but when accounting for transfection rate, the overall knock-in efficiency (0.35/16 = 2.1%) is consistent with experiments reported above (Supplemental Fig. S5).

**Figure 8.**
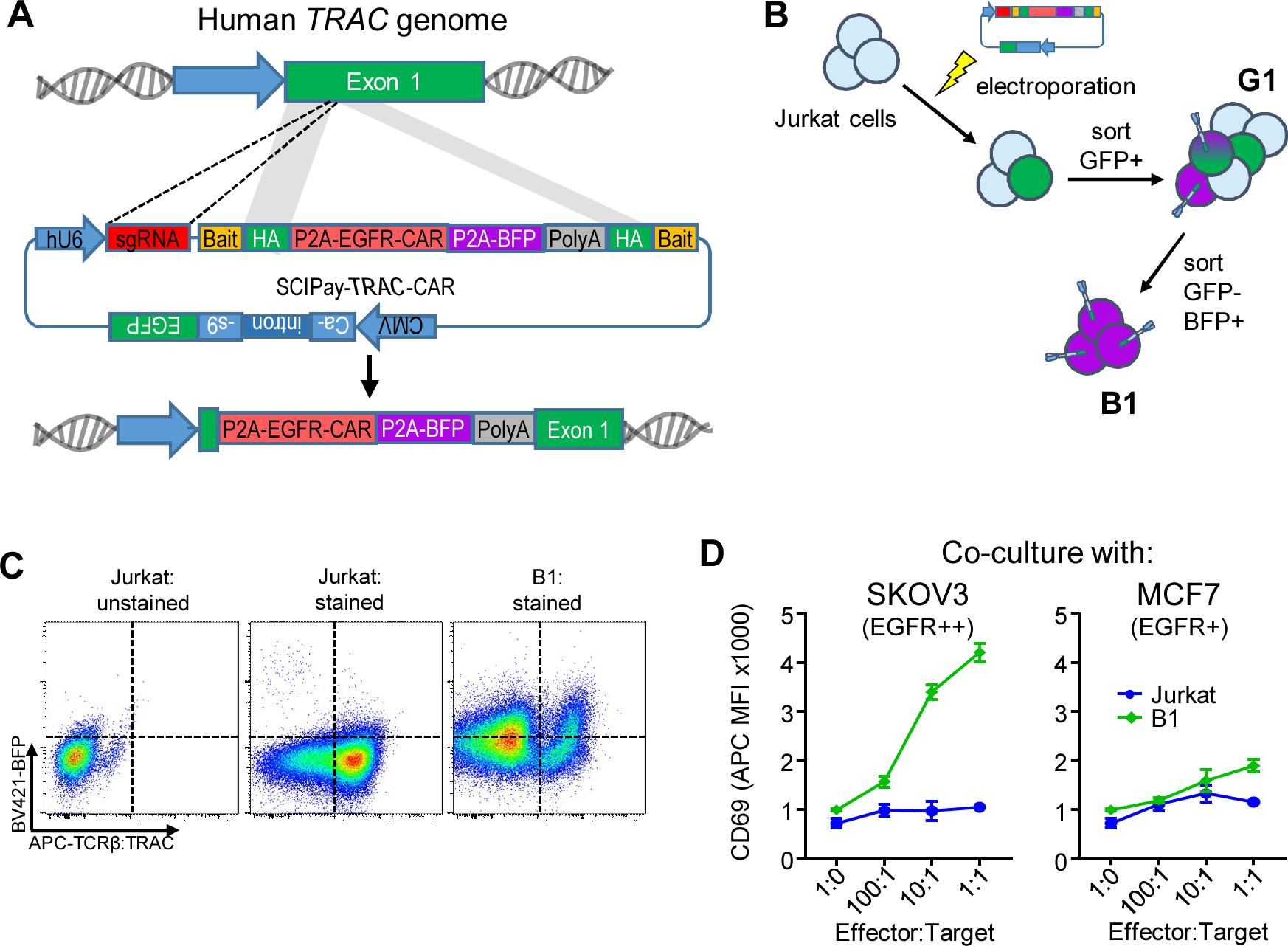
Using SCIPpay to insert a chimeric antigen receptor (CAR) into the T cell receptor alpha subunit (*TRAC*) gene. (A) Overview of SCIPpay designed to insert an anti-EGFR CAR with BFP into human *TRAC*. (B) Outline of cell engineering strategy. After electroporation, successfully transfected cells were sorted twice to enrich BFP-positive cells. (C) BFP and TCR/*TRAC* expression on double-sorted population (B1). (D) Surface CD69 expression during co-culture between the indicated cell population and SKOV3 or MCF7 cells. Results in (D) represent means +/- SEM of 3 independent experiments.

Two subsequent sorting rounds yielded a population (B1) that was 40% BFP-positive and 90% TRAC/TCRβ-negative (Fig. 8C). Notably, we consistently observe that Jurkat cells cycle between 50-80% TCR positivity (see middle panel of Fig. 8C) and here, the CAR-BFP is driven by *TRAC* and not a constitutive promoter. Therefore, 40% is likely under-representative of the total CAR-positive cells in B1. Regardless, when co-cultured alongside two cancer cell lines with different EGFR expression (SKOV3, EGFR++; MCF7, EGFR+), B1 increased surface CD69 expression in response to increasing target cell ratios, while normal Jurkat cells did not (Fig. 8D). This response also stratified by EGFR expression on target cells, as SKOV3 caused greater CD69 induction than MCF7 (Fig. 8D). Overall, these results demonstrate that SCIPpay is an effective engineering tool.

## Discussion

Numerous strategies for genome editing have been conceived, with several showing promise as scalable technologies with therapeutic intentions. Although many demonstrate robust function, these protocols typically employ relatively complex technical processes, including RNA, protein, and/or virus manufacturing and handling. Here, we tested two relatively simple and cost-effective self-cleaving plasmid formats for their ability to perform gene editing, which enabled the delivery of all necessary components using a single DNA molecule without requiring virus/RNA/protein production.Several SCIP-like technologies have appeared while performing the work presented here. In particular, Anja Ehrhardt’s team at Witten/Herdecke University employ AdV/AAV for delivering HDR-compatible DNA editing machinery (Ehrke-Schulz et al. 2017; Gao et al. 2018; Bergmann et al. 2018; Schiwon et al. 2018).

While previous editing strategies are robust and predictable when optimized, they all involve more than one component, with most also requiring RNA, protein, and/or virus production. These entities are more difficult and complex to generate and handle than plasmids). Instead, SCIPs incorporate all the elements required for target-specific genome engineering any other helper component. For simple knock-out cell line generation, SCIPs are created by two sequential or simultaneous restriction/ligation reactions with short annealed oligonucleotides, meaning novel SCIPs can be generated in as few as 2 days with little hands-on time. Once produced, SCIPs can be immediately transfected into cells which are then assessed for functional effects, where according to our observations 50-80% of stable SCIP integrant cells should be complete knockout. Additionally, as a standard SCIP integration results in permanent Cas9 expression, this may potentially simplify subsequent serial editing strategies. While constructing SCIPpays for genomic knock-in of novel transgenic payloads is slightly more complicated and costly, involving higher-level genetic design and DNA fragment synthesis, the process is still very straightforward and should be achievable by any lab with basic molecular biology capabilities.

HDR-mediated genome editing success rates with SCIPpay were relatively modest (3% with 500 bp HAs), although this was more than adequate to engineer useful cell lines, and further refinement to the genetic design and delivery protocol are ongoing to optimize these for specific purposes. While other nuclease-based systems have demonstrated higher efficiencies, these protocols are more complex and have different applications than SCIPpay. In fact, we believe that SCIPpay represents the easiest and most accessible method to simultaneously cause knock-in and knock-out, as illustrated by inserting a tdTomato transgene into the CD69 locus, or CAR into *TRAC*, if such an outcome is desired. Furthermore, SCIPpay can be easily adapted to screen the effects of delivering the same transgenic payload to a different of genomic sites, for high-throughput optimization of therapeutic gene editing approaches. SCIPs can also be adapted to enable iterative genome editing, as this process is non-viral and there is less chance for cells to build up immunity/resistance to subsequent engineering efforts. Another key advantage relating to the single-plasmid nature of SCIPs, is that they potentially permit the creation of integrating plasmid arrays and/or libraries which could be readily applied to identifying optimal sites for transgene delivery. An additional tantalizing application we suggest is using SCIP libraries to identify novel safe harbor sites in various genomes.

Our data do not permit definitive conclusions regarding SCIP’s integration mechanism (NHEJ, microhomology, or homology-directed repair mediated), as the NHEJ orientation without micro-homology is selected against due to re-creating the gRNA target sites (Supplemental Fig. S2B). We do note that expanding homology to 500bp resulted in only a 2-fold increase in knock-in efficiency supporting a emerging view that microhomology or homology-independent mechanisms can be used for predictable transgenic integration. Most relevant to SCIP, these results call into question whether the added complexity of an HDR integration strategy is justified; instead use of SCIP or SCIPpay with minimal bait sites and a robust selection strategy seems to be the easiest means of isolating desired clonal integrants. We also noted a surprisingly high percentage of SCIP knock-in to unknown loci (precision scores never exceeded 80%) and that 14% of PCR-tested knock-out plus knock-in clonal cell lines did not possess apparent on-target SCIP integration (Table 1). We hypothesize that this is potentially gRNA-dependent and indicative of gRNA promiscuity; in fact, this may represent a valuable assay for measurement and optimization of such attributes. We are now testing a large number of SCIP plasmids across a range of surface expressed gene targets to assess whether SCIP can be used to quickly assess on- and off-target cutting rates.

We note that other nuclease-enabled gene editing technologies have demonstrated targeted integration of DNA payloads up to 15 kb (Maresca et al. 2013), more than we have shown here with SCIPpay. While molecular biology dogma suggests that smaller plasmids are easier to work with, a 15 kb payload could be achieved with a SCIPpay of 20 kb (500 bp HAs and keeping the T2A-EGFP following Cas9). Using the commonly-used AdV cloning plasmid AdEasy which is 33.5 kb in size, a SCIP plasmid could deliver a 24 kb payload. While still hypothetical, we feel SCIP may offer the most accessible means to perform targeted delivery of large transgenic cargo to genomic targets.

Lastly, alongside SCIP’s benefits as a modular DNA editing tool are potential safety considerations. Although genetic transfer via plasmids should be less efficient than viruses, SCIPs act essentially as selfish genetic elements and thus we chose to treat SCIPs as BSL2 biohazards during their cloning and purification. Despite this precaution, depending on the transgenic sequences being delivered we do not believe that SCIPs pose general safety risks in excess of other gene editing tools.

## Conclusion

We present here development and proof of concept application of a novel gene editing strategy based on self-cutting and integrating plasmids (SCIPs). We conclude that SCIPs represent an extremely cost-effective and accessible strategy for delivery of novel transgenic DNA to targeted genomic sites, and can be employed to create useful engineered cell lines. In addition to the uses demonstrated here (ie. making knock-out cell lines and inserting DNA payloads at targeted sites), our SCIP system shows promise in advanced and unique situations, such as measuring and optimizing on- and off-target integration efficiency, safe harbor site identification, and performing library-scale multiplexed and/or iterative automated genome engineering. Therefore, we believe that SCIPs represent innovative technologies and intend that their presentation here will stimulate their use by the wider research community.

## Methods

### Cell Culture

Jurkat, EL4, Raji (ATCC) and Nalm6 (generously provided by Dr. Beat Bornhauser, University Children’s Hospital Zurich) cells were cultured in standard RPMI media supplemented with 10% fetal bovine serum (FBS), 1% penicillin/streptomycin, 1 mM sodium pyruvate, 2 mM L-glutamine, and 55 μM β-mercatpoethanol at 37°C and 5% CO2. HEK293T, SKOV3, and MCF7 cells (ATCC) were cultured in similar DMEM-based media. Cell counting was performed using an automated cell counter (Cellometer; Nexcelcom, Lawrence, MA) which assesses live/dead counts using acridine orange/propidium iodide staining.

### Plasmids

The Cas9-CRISPR plasmid pSpCas9(BB)-2A-GFP (px458) (Addgene #48138) (Ran et al. 2013) served as the source of pSCIP development. The pQC (“quick-change”) backbone plasmid containing a second golden gate cloning site for CRISPR bait insertion was created by cloning synthetic DNA oligonucleotides (Alpha DNA, Montreal, Canada) encoding tandem oppositional BsaI sites between the XbaI and KpnI sites of px458 (Supplemental Fig. S1A). Similar to the existing BpiI-based golden gate cloning cassette in px458 for gRNA DNA insertion, this enables parallel or serial bait DNA insertion via BsaI-based Golden Gate assembly. An intron derived from the mouse immunoglobulin precursor V-region (GenBank ID: AH002574.2) was then inserted via Gibson cloning (300 ng digested plasmid and 10 ng DNA fragments with 1x in-house assembly mixture) into the middle of the SpCas9 coding sequence, as previously demonstrated to limit leaky SpCas9 expression in bacteria (Petris et al. 2017). This plasmid is denoted pQCi (quick change with an intron). The following plasmids reported in this manuscript have been deposited in the Addgene plasmid repository: pQCi (Addgene #154086), pQCi-mB2m-sgRNA (Addgene #154090), pSCIP-mB2M (Addgene #150091), pQCi-hTRAC-sgRNA (Addgene #154093), pSCIP-hTRAC (Addgene #154094), pSCIPpay-hCD69-TdTomato (Addgene #154095), pSCIPpay-BFP reporter (Addgene #154096), and pSCIPpay-BFP reporter-mB2M-500Has (Addgene #154097).

CRISPR guides were designed using CRISPOR (http://crispor.tefor.net/) and potential sequences were curated manually (targeting early exons, high predicted efficiency and specificity, low/non-coding off-targets) (Supplemental Table S1). Corresponding complimentary DNA oligonucleotides were then synthesized (Alpha DNA, Montreal, Canada) and cloned using a one-step restriction/ligation reaction involving 300 ng plasmid DNA, 1 µM annealed oligonucleotides, 0.25 µL/2.5 U BpiI (ThermoFisher ER1012), 0.5 uL/0.5 U T4 ligase (ThermoFisher 15224017), and T4 ligase buffer (ThermoFisher 46300018). To create self-cutting plasmids, corresponding bait DNA sequences including PAM sites were inserted using a similar protocol involving 0.25 µL/2.5 U BsaI (ThermoFisher ER0291) (Supplemental Fig. S1A). These restriction/ligation reactions involving annealed oligonucleotides are extremely efficient and result in successful cloning in greater than 95% of antibiotic-resistant clones.

SCIPs targeting mouse *B2m* with varying HA lengths and orientations were created using a similar approach (Supplemental Fig. S1B). For inverted HAs, synthetic DNA fragments of the mouse *B2m* gene (GenBank ID: 12010, assembly GRCm38.p6) were constructed (Twist Bioscience, San Francisco, CA) in defined lengths surrounding the CRISPR target site (Supplemental Table S1) with capping sequences allowing their seamless cloning via BsaI-based restriction/ligation. For forward HAs, synthetic DNA fragments were similarly constructed (Twist Bioscience) with BsaI cloning caps such that the *B2m* genome sequences (not their reverse complements) on either side of the CRISPR site were exchanged for each other, thereby providing “normal” homology upon plasmid linearization (see Fig. 5A). These reactions consisted of 300 ng plasmid DNA, 10 ng DNA fragments, 0.25 µL/2.5 U BsaI (ThermoFisher, Waltham, MA; ER0291), 0.5 uL/0.5 U T4 ligase (ThermoFisher 15224017), and T4 ligase buffer (ThermoFisher 46300018). Note that forward homology plasmids contain 17 or 6 base pair non-homologous caps resulting from Cas9 cleavage of the bait site.

To create the pSCIPpay-BFP-reporter backbone plasmid, a DNA sequence was constructed *in silico* containing a P2A-BFP transgene without BpiI, BsaI, and BsmBI restriction sites (subsequently validated to maintain functionality) containing terminal BsaI cloning cassettes with distinct cloning sticky ends (Supplemental Fig. S1C). This DNA fragment (Twist Bioscience) was inserted into BsaI-digested pQCi using Gibson assembly. This modular plasmid allows insertion of gRNA-coding DNA oligonucleotides using BpiI as outlined above and subsequent cloning of HA- and bait-coding DNA fragments engineered to be in-frame with the preceding genome sequence and following P2A-BFP sequence with BsaI using Gibson or restriction/ligation cloning.

SCIPpay-human-*CD69*-tdTomato and SCIPpay-human-*TRAC*-CAR were generated by first cloning gRNA-coding DNA oligonucleotides as described above and subsequently inserting DNA fragments (Twist Bioscience) designed *in silico* with gRNA-specific bait sequences into pQCi using Gibson assembly. EGFR-targeting CAR utilized a novel EGFR-targeting sequence under development at the National Research Council Canada.

Plasmid DNA was purified after overnight culture in standard lysogeny broth containing 100 µg/mL ampicillin using a miniprep plasmid DNA isolation kit (Sigma Aldrich, St Louis, MO; PLN350) and quantified using spectrophotometry (NanoDrop One, ThermoFisher). Plasmid construction was always validated to *in silico* sequences using Sanger sequencing.

### Transformation and Colony Counts

Standard *Escherichia coli* DH5α chemically competent cells generated using an in-house protocol from an in-house stock were used for most experiments. Stbl3 (ThermoFisher) and BL21 (New England Biolabs, Ipswich, MA) cells were utilized when indicated. Briefly, 50 µL freshly thawed cells were incubated on ice with 10 ng plasmid DNA for 30 minutes, heat shocked at 42°C for 35 seconds, and placed on ice for 2 minutes. Cells were recovered by adding 950 µL warmed SOC media and incubating at 37°C while shaking vigorously for 1 hour. From this solution, 100 µL was plated on ampicillin-containing agar plates and incubated overnight at 37°C. Colony counts were performed the following morning and reported values represent actual numbers of colonies, not normalized for dilutions.

### Transfection

EL4 and Jurkat cells were transfected via electroporation according to a previously-outlined protocol (Chicaybam et al. 2013). After collection, 5×10^5^ – 1×10^6^ cells were suspended in 100 μL Buffer 1SM (5 mM KCl, 15 mM MgCl2 120 mM Na2HPO4/NaH2PO4, 25 mM sodium succinate, and 25 mM mannitol; pH 7.2) and briefly (<5 minutes) incubated with appropriate plasmid DNA (2-5 μg). This solution was transferred into 0.2 cm generic electroporation cuvettes (Biorad Gene Pulser; Bio-Rad laboratories, Hercules, CA) and immediately electroporated using a Lonza Nucleofector I (Lonza, Basel, Switzerland) and program X-05 (X-005 on newer Nucleofector models). Cells were cultured in pre-warmed recovery media (RPMI with 20% FBS, 2 mM L-glutamine and 1 mM sodium pyruvate) overnight before being transferred to normal culture media.

### Flow Cytometry

Transfection, Knock-out, and Knock-in Detection Transfection efficiencies were evaluated on cells 24 and 48 hours post-transfection. Cells were collected from culture, centrifuged for 5 minutes at 300g, re-suspended in flow cytometry staining buffer (FSB, PBS with 1% FBS, 10 mM HEPES, and 2 mM EDTA), and analyzed for GFP using the FITC channel.

PDCD1 and B2M expression on EL4 cells were assessed with fluorescent antibody staining using an APC-conjugated PD1 antibody (BioLegend, San Diego, CA: 135210) or AlexaFluor 647-conjugated H-2K^b^ antibody (BD Biosciences, San Jose, CA; 562832). Briefly, cells were collected from culture, centrifuged at 300g for 5 minutes, re-suspended in antibody diluted in FSB (2.5 µL per million cells), incubated at room temperature for 30 minutes, centrifuged at 300g for 5 minutes, re-suspended in flow buffer, fixed with 0.5% formalin in PBS, and immediately analyzed. TCR expression on Jurkat cells was similarly assessed using a BV510-conjugated TCRαβ antibody (BD Bioscience 563625) or APC-conjugated CD3 antibody (BD Bioscience 561810).

As SCIP integration results in CMV-driven Cas9-GFP expression, knock-in events were detectable by simultaneous assessment of FITC fluorescence. Similarly, SCIPpay-BFP-reporter and SCIPpay-*TRAC*- CAR knock-in was assessed using the BV421 filter set and SCIPpay-CD69-tDtomato was assessed using the PE filter set.

#### Jurkat Functional Testing

Jurkat activation during anti-CD3 administration or co-culture was assessed using flow cytometry detection of CD69 with an APC-conjugated anti-CD69 antibody (BD Bioscience 555533). Cells were not washed prior to or after staining: 0.2 μL antibody was added to 50 µL PBS per well of a 96-well plate, this diluted antibody solution was added to wells, cells were incubated for 30 min at 37°C, formaldehyde was added to a final concentration of 0.5%, and the plate was immediately analyzed. For TRAC-CAR knock-in, Jurkat cells were distinguished from target cells via BFP (BV421).

Jurkat-CD69-tdTomato (J69) cell activation was assessed by detecting PE fluorescence and for J69-CAR, Jurkat cells were distinguished from target cells via tdTomato (PE) and GFP (FITC).

All flow cytometry analyses were performed using a BD LSR Fortessa configured with a high-throughput plate reader and the following lasers: 355 nm, 405 nm, 488 nm, 561 nm, and 640 nm.

#### Cell Line Generation Using SCIPs

Stable cell lines (clonal EL4-*B2m*-SCIP, J69, Jurkat-*TRAC*-CAR) were generated using high-speed fluorescence-based cell sorting using various staining and gating strategies as described in the text (MoFlo Astrios, Beckman Coulter, Mississauga, ON).

### Genomic DNA Extraction, PCR, and Sequencing

Crude genomic DNA extracts were generated from clonal EL4 cells to confirm *B2m*-SCIP integration. Briefly, cell clones expanded to 50-100,000 cells were centrifuged, re-suspended in 100 µL lysis buffer (50 mM NaOH, 0.2 mM EDTA), incubated at 98°C for 15 min, cooled to 4°C, and 10 µL neutralization buffer (1 M Tris-HCl, pH 7.0) was added to each sample. DNA was then frozen at -80°C for short-term storage.

PCR was performed using DreamTaq Green (ThermoFisher K1082) and combinations of the primers indicated in Table S3 to detect SCIP integration orientation and efficiency. Products from select reactions were then Sanger sequenced using either U6-F, CMVe-R, or *B2m*-R primers (Table S2).

### Lentivirus Production and Jurkat Transduction

CD19-CAR lentivirus was generated using HEK293T cells. Briefly, 400,000 HEK cells were seeded in 6-well plates and the next day transfected with 1 μg psPAX2 (Addgene #12260), 3 μg pCMV-VSV.G (Addgene, Watertown, MA; #8454), and 4 μg CD19-CAR-GFP transfer plasmid (Bloemberg et al. 2020) with 24 μg linear PEI (Polysciences Inc., Warrington, PA) for 6 hours. Crude culture media supernatants containing virus particles were collected 72 hours later, centrifuged at 500g for 10 min at 4°C to remove cell debris, and filtered through a 0.45 μm PVDF filter (Millipore/Sigma).

To make CAR-expressing Jurkat-CD69-reporter cells, 250,000 cells were incubated with a 1:1 mixture of this filtered viral supernatant and normal culture media containing 10 μg/mL Dextran for 48 hours. Cells were then washed and expanded in normal culture media. As the CAR transgene is followed by a P2A-GFP construct in the lentiviral transfer plasmid, all CAR-positive cells express GFP. Therefore, resting J69-CAR19 cells are CD69-negative, faintly tdTomato/PE-positive, and GFP-positive. Upon activation by co-culture with CD19 positive cells; these J69-CAR cells exhibit increased expression of tdTomato/PE.

### Continuous Live-Cell Imaging

Continuous live-cell imaging (IncuCyte® S3, Sartorius) was used to assess J69 activation. After adding anti-CD3 antibody or CAR target cells, images were acquired every 30-60 minutes in light/phase and red (ex. 565-605 nm; em. 625-705 nm) fluorescence channels.

## Statistics

Quantified results presented are means +/- standard error (SEM) and typically represent 3-5 independently run experiments; specific experiment descriptions are contained in respective figure captions.

Statistical comparisons were performed in appropriate situations where group size and replication numbers were not predetermined for power effects. T tests were calculated using Microsoft Excel and ANOVA analyses were performed using GraphPad Prism. Tukey’s multiple comparison post hoc was used to identify locations of significance for ANOVAs. A p-value of 0.05 was considered statistically significant in all situations. Analysis specifics and descriptions are included in corresponding figure captions.

## Data Access

All data generated in the course of experiments reported here are available upon request. Sequencing data files and plasmid maps for plasmids used have been provided here as supplemental material and can be obtained from the Addgene repository, with the exception of pSCIPpay-TRAC-CAR which can be obtained under MTA.

## Acknowledgements

The authors wish to thank the National Research Council Canada’s genomics core facility, including Dr Qing Yan Liu and Joy Lei for technical expertise and advice for performing Sanger sequencing runs. This work was funded by the National Research Council Canada Disruptive Technology Solutions Cell and Gene Therapy challenge program.

## Author Contributions

SCIP conception (SM, DB), study design and planning (all), conducting experiments and collecting data (DB, DSM, SM, TN), data analysis and figure preparation (DB, DSM, SM), and manuscript writing (DB, SM). All authors approved of the final version of this manuscript and figures prior to its submission.

## Supplemental Material

**Supplemental Figure S1.**
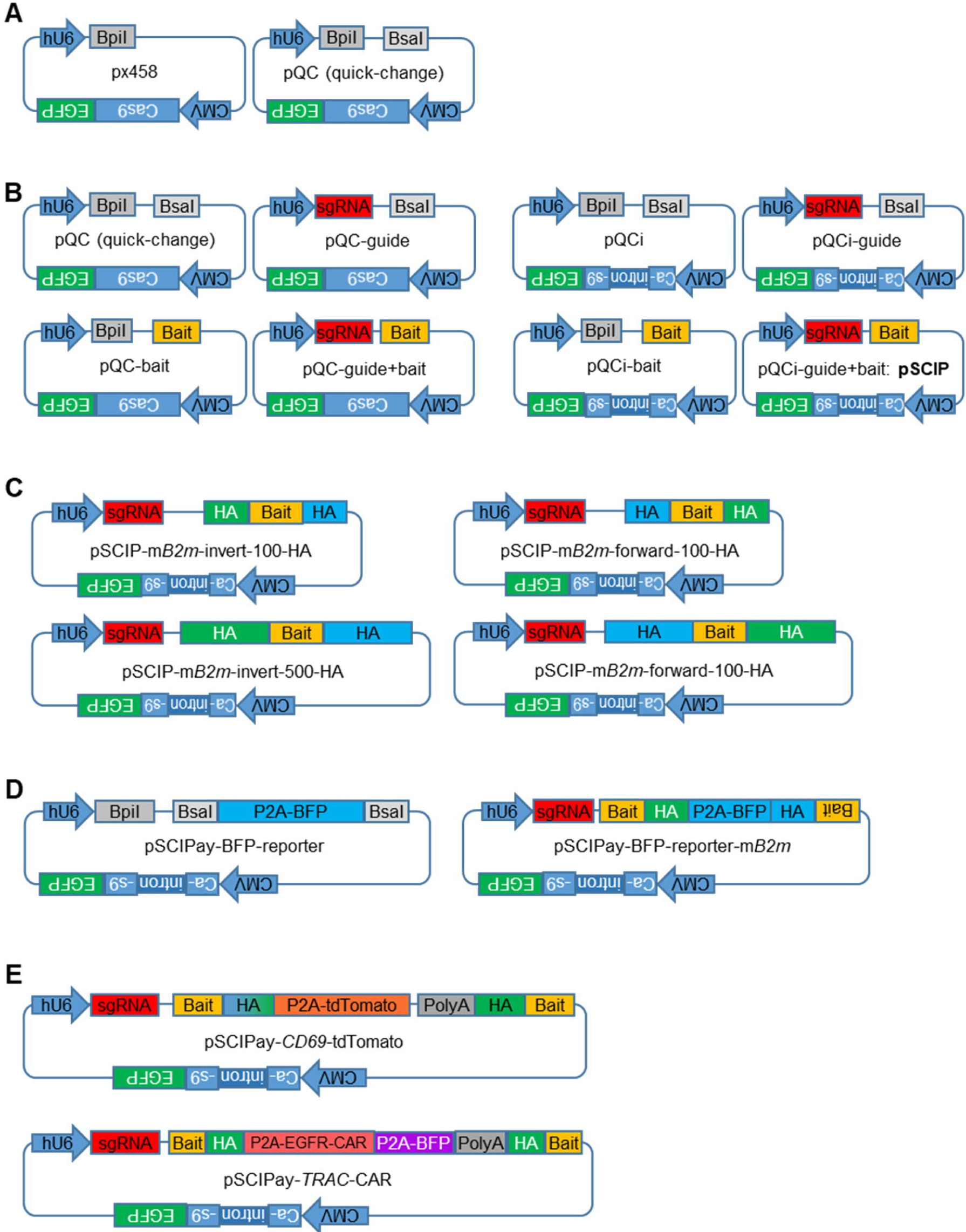
Overview and nomenclature of relevant plasmids. (A) The widely used Cas9-GFP plasmid px458 served as the backbone for SCIP development, which began by inserting a second “golden-gate” restriction site (BsaI) for inserting bait sequences alongside the sgRNA cloning site (BpiI). (B) Combinations of guide- and/or bait-containing plasmids without (left side) or with (right side) the mammalian intron as used in Figures 2-4. Final “pSCIP” structure is pQCi-guide+bait: pQC with the intron, with the sgRNA, and with the bait sequence. (C) Structural depictions of pSCIPs with varying homology arm lengths and orientations, as used in Figure 5. (D) Structure of pSCIPay-BFP backbone/cloning plasmid, along with mouse B2M-targeting version, as used in Figure 6. (E) Structure of SCIPays used for cellular engineering in Figures 7 and 8.

**Supplemental Figure S2.**
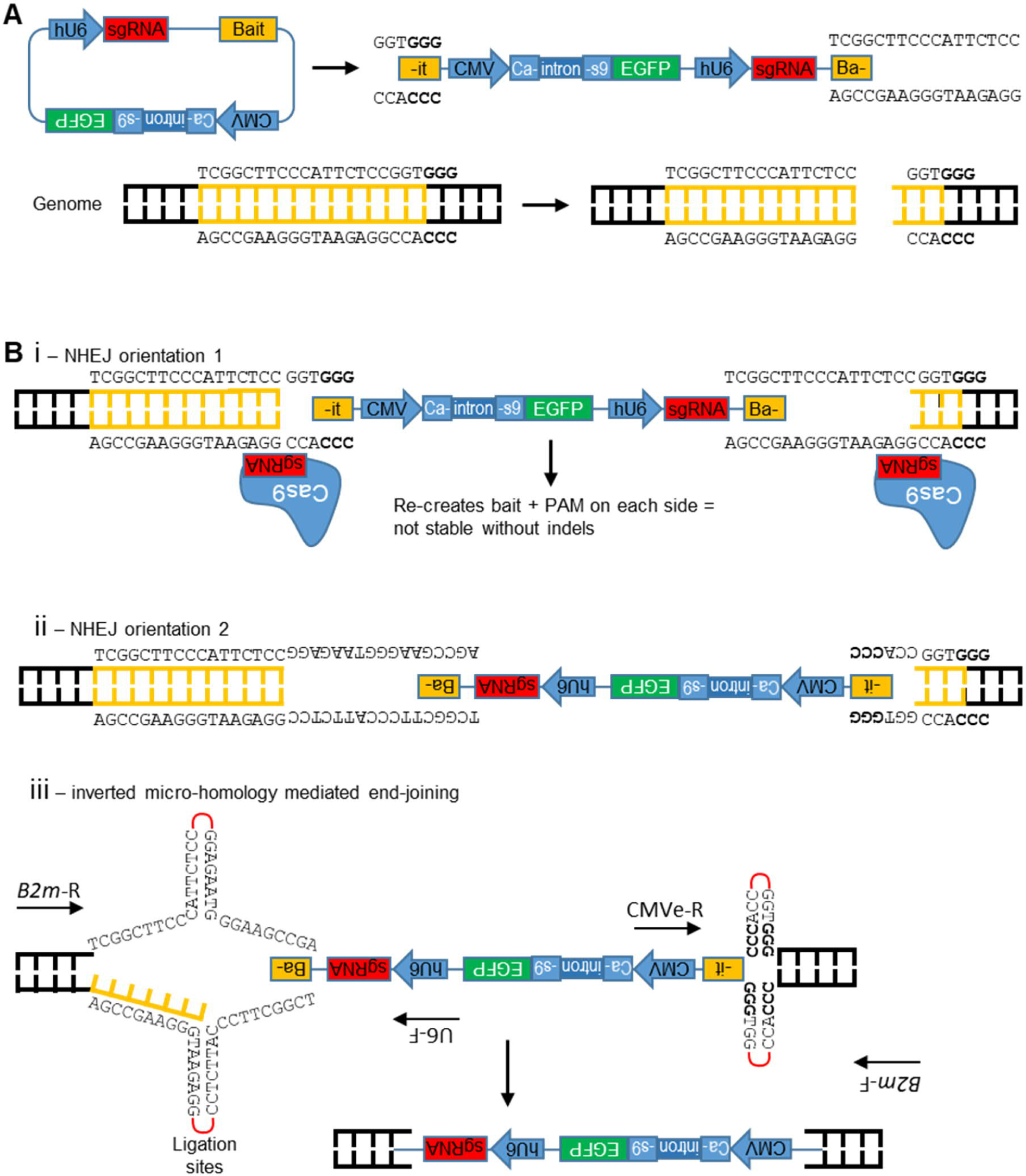
SCIP genomic integration mechanism possibilities. (A) Upon Cas9/sgRNA production and RNP formation, DSBs are made in genomic (bottom) and plasmid (top) DNA sequences, producing the indicated genetic structures. This example illustrates the mouse *B2m* site/gRNA used throughout the manuscript. (B) Potential integration mechanisms. (Bi) NHEJ orientation 1 re-creates the Cas9/gRNA site and should be selected against. (Bii) NHEJ in the opposite orientation leads to SCIP integration and gene interruption. (Biii) Due to CRISPR guide and bait structures, short sequences of inverted homology exist for NHEJ orientation 2, potentially contributing to site- and direction-specific integration. Figure 4 and Table 1 demonstrate this is in fact true.

**Supplemental Figure S3.**
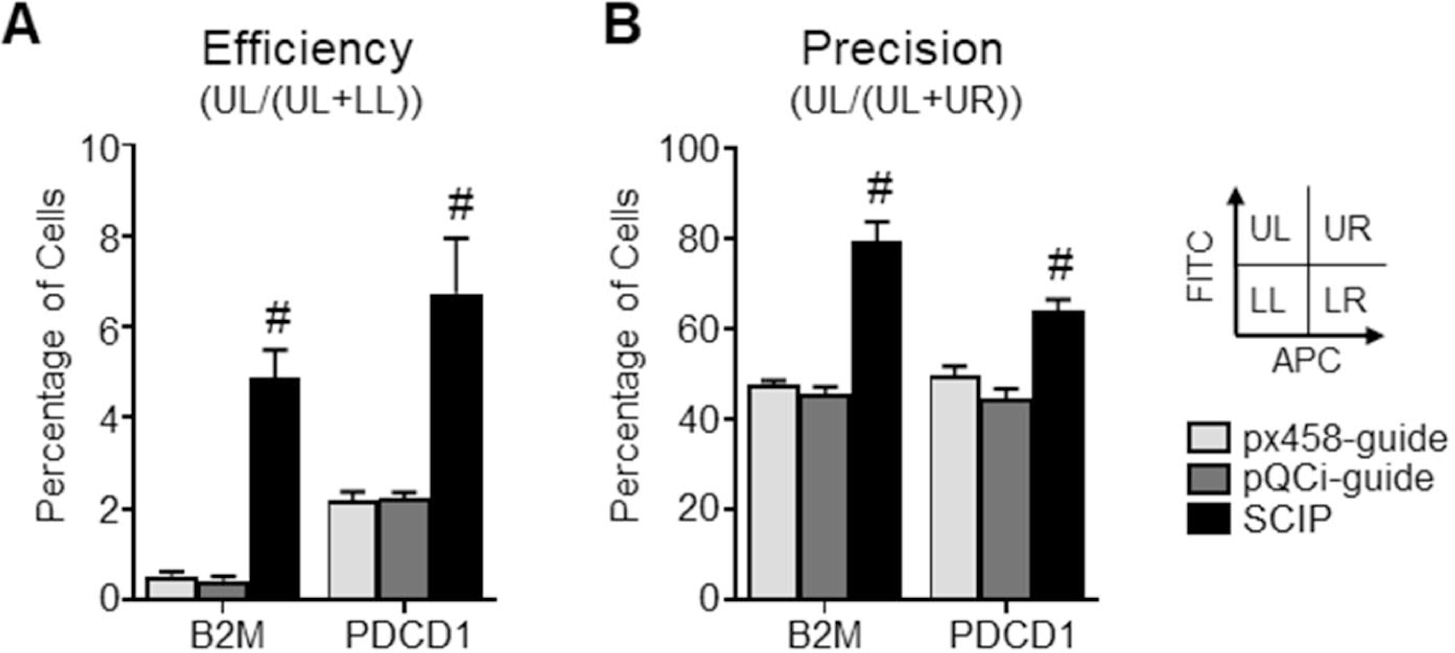
SCIP integration 14 days post-transfection. This data supplements Figure 4B, where EL4 cells were electroporated with the indicated plasmids and assessed for gene knock-out and SCIP knock-in 14 days later. (A) Efficiency and (B) precision scores, similar to those presented in Figure 4B. Results represent means +/- SEM of 4 independent experiments. Pound signs (#) represent significant difference between the indicated group and all other groups, as calculated using 1-way ANOVA (p<0.05).

**Supplemental Figure S4.**
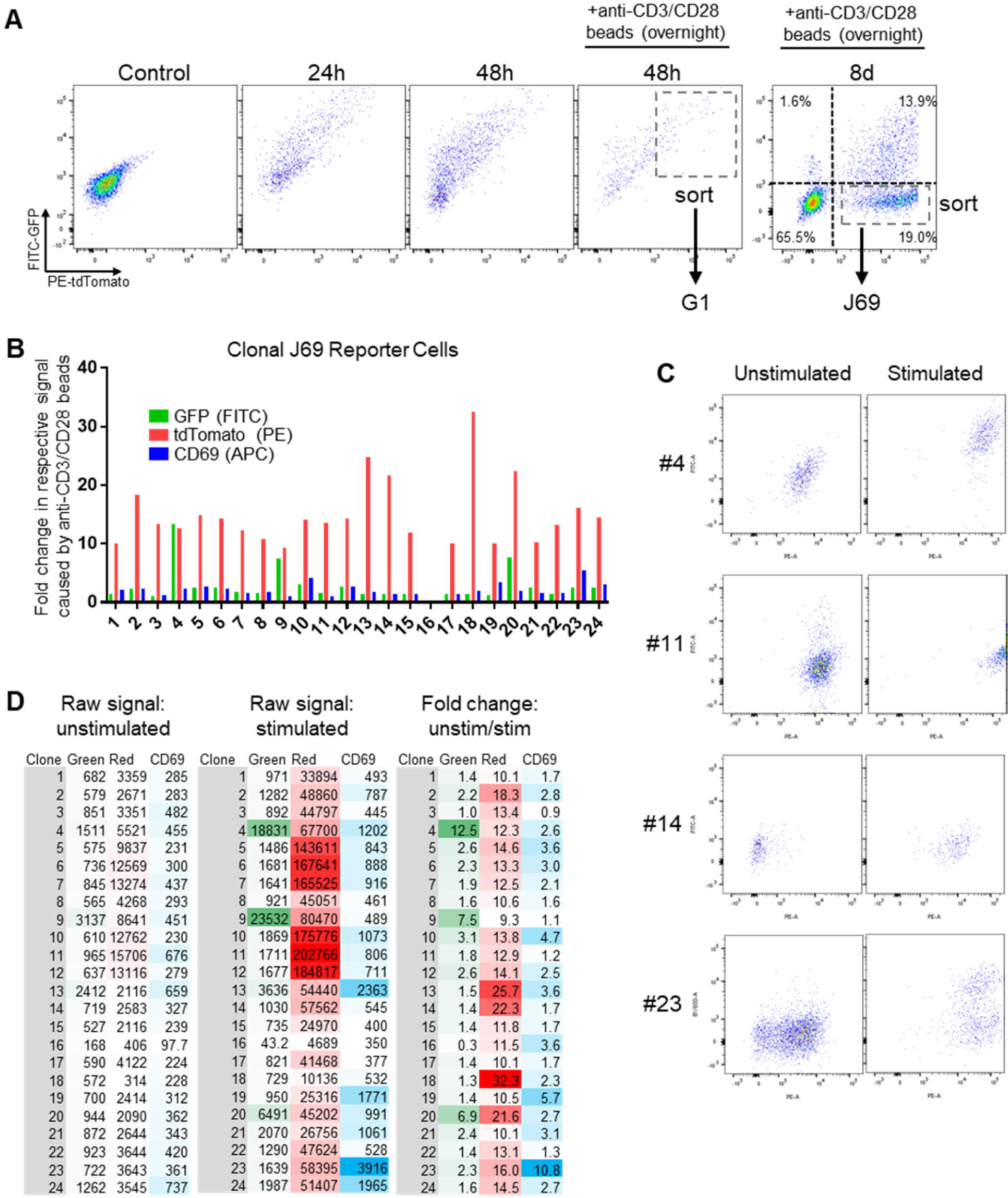
Generating clonal J69 cell lines. To derive clonal J69 cells, the polyclonal population was single-cell sorted and 24 surviving clones were assessed for T cell responses. (A) Overview of flow cytometry gating and sorting analyses. Clonal cells were sorted from the J69 populations (B) FITC-GFP, PE-tdTomato, and CD69-APC responses after incubating cells with anti-CD3/CD28 beads. Data are expressed relative to non-stimulated cells. (C) Representative flow cytometry plots from select clones. (D) Raw green (FITC-GFP), red (PE-tdTomato), and CD69 (APC) mean fluorescence intensities.

**Supplemental Figure S5.**
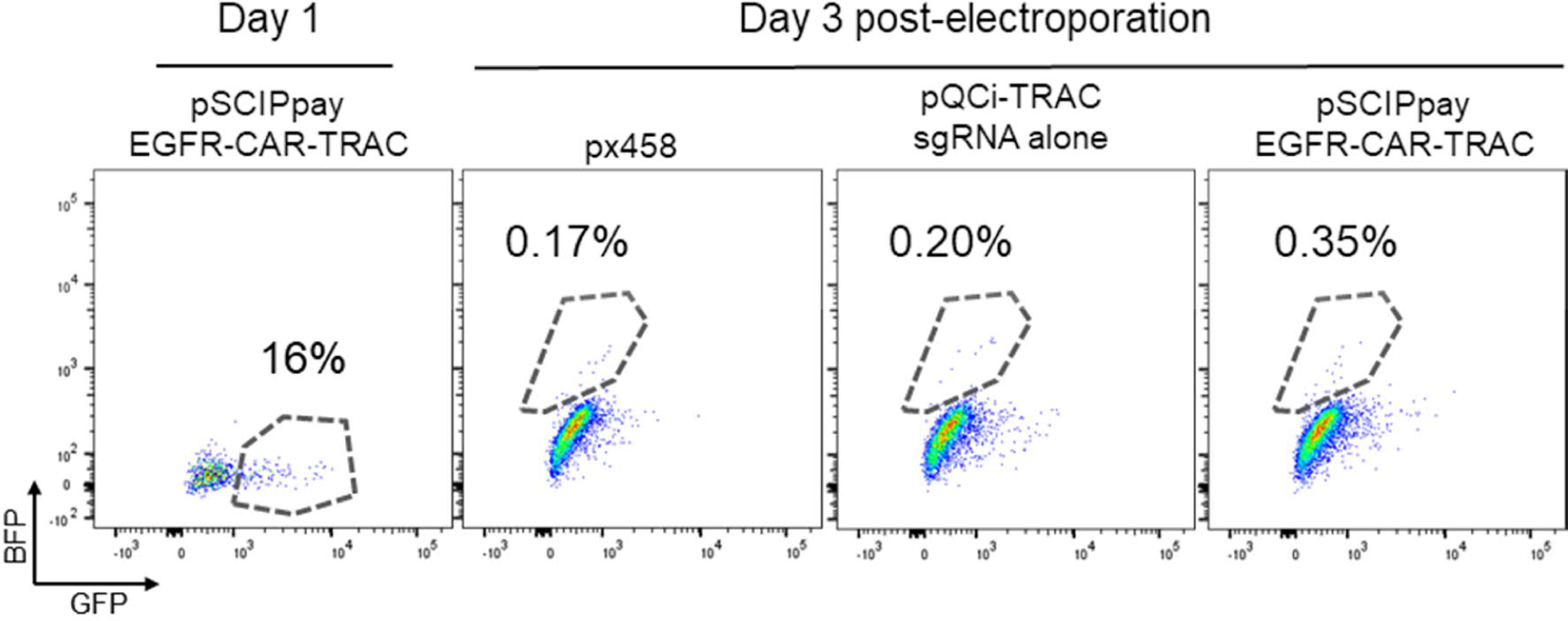
Flow cytometry gating for Figure 8.

**Supplemental Table S1:**
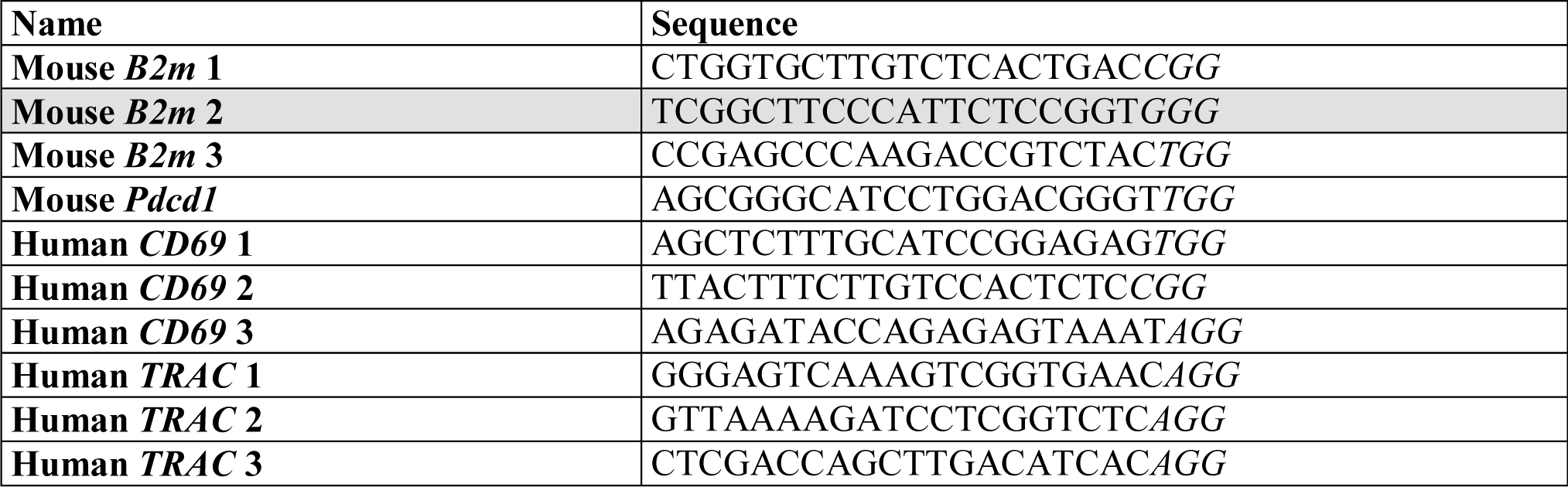
CRISPR gRNA Target Sequences.

**Supplemental Table S2:** EL4-*B2m*-SCIP Genomic PCR Possibilities.

| Primer Pair | Size (bp) | #1 | #2 | #3 | #4 | #5 | #6 |
| --- | --- | --- | --- | --- | --- | --- | --- |
| *B2m*-F + *B2m*-R | 345 (WT) | No | No | No | Maybe/<br>probably | Maybe/<br>probably | Maybe/<br>probably |
| U6-F + *B2m*-R (predicted) | 558 | Yes | No | Yes | Yes | No | No |
| U6-F + *B2m*-F (backwards) | 541 | No | Yes | Yes | No | Yes | No |
| CMVe-R + *B2m*-F (predicted) | 410 | Yes | No | Yes | Yes | No | No |
| CMVe-R + *B2m*-R (backwards) | 428 | No | Yes | Yes | No | Yes | No |

**Supplemental Table S3:**
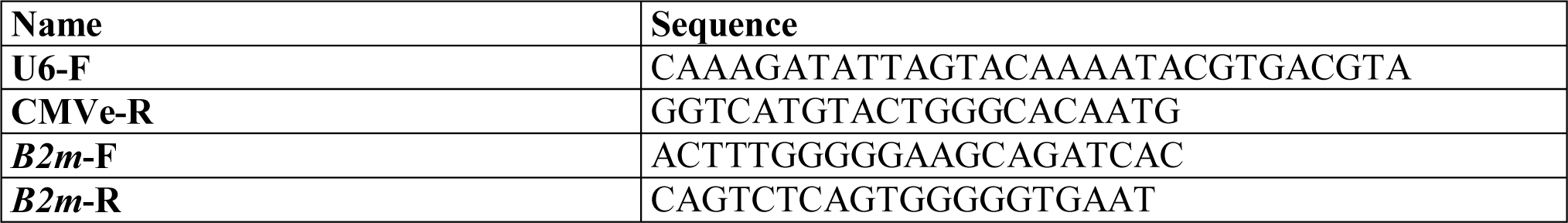
PCR Primers.

### Supplemental Sequencing Data Description

We have included raw Sanger files (in .ab1 format) that correspond to Figure 4F where we sequenced the SCIP integration “seams” at the *B2m* locus in clonal EL4 cells. These files are labelled with the letter “C” or “U” along with a number, whereby:

C – indicates the PCR product was sequenced with our CMVe-R primer (Supplemental Table S3)

U – indicates the PCR product was sequenced with our U6-F primer (Supplemental Table S3)

\# - represents the clone number, according to Figure 4E and Table 1

We include a file containing the predicted sequence where the entire SCIP plasmid integrates into the corresponding CRISPR target site within the murine B2M gene. Genbank format (.gb) sequence map files are also provided for plasmids that have been deposited in the Addgene plasmid repository: pQCi (Addgene #154086), pQCi-mB2m-sgRNA (Addgene #154090), pSCIP-mB2M (Addgene #150091), pQCi-hTRAC-sgRNA (Addgene #154093), pSCIP-hTRAC (Addgene #154094), pSCIPpay-hCD69-TdTomato (Addgene #154095), pSCIPpay-BFP reporter (Addgene #154096), and pSCIPpay-BFP reporter-mB2M-500Has (Addgene #154097).

